# Estimating tumor mutational burden from RNA-sequencing without a matched-normal sample

**DOI:** 10.1101/2021.08.15.456379

**Authors:** Rotem Katzir, Keren Yizhak

## Abstract

Detection of somatic point mutations using patients sequencing data has many clinical applications, including the identification of cancer driver genes, detection of mutational signatures, and estimation of tumor mutational burden (TMB). In a recent work we developed a tool for detection of somatic mutations using tumor RNA and matched-normal DNA. Here, we further extend it to detect somatic mutations from RNA sequencing data without a matched-normal sample. This is accomplished via a machine learning approach that classifies mutations as either somatic or germline based on various features. When applied to RNA-sequencing of >450 melanoma samples high precision and recall are achieved, and both mutational signatures and driver genes are correctly identified. Finally, we show that RNA-based TMB is significantly associated with patient survival, with similar or superior performance to DNA-based TMB. Our pipeline can be utilized in many future applications, analyzing novel and existing datasets where only RNA is available.

## Introduction

Somatic point mutations accumulate in the DNA of all dividing cells, both normal and neoplastic, and are the most common mechanism for altering gene function [1]–[4]. Their detection in tumor samples is of high clinical value; first, when accumulated in specific genes termed “drivers", they may lead to the development of cancer. Identifying these mutations is therefore crucial for matching existing targeted therapies and for developing novel ones [5]–[8]. In addition, somatic mutations are used to determine intra-tumor heterogeneity which is a major mechanism of therapeutic resistance [9], and for identifying mutational signatures that have proven as clinically useful biomarkers [10], [11]. More recently, the set of somatic mutations in a tumor has been used to estimate the tumor mutational burden (TMB), an emerging proxy for neoantigen load. TMB is defined as the number of non-silent mutations found in a tumor DNA, and was found to be an independent marker of patient response to immune checkpoint inhibitor therapy (ICI), and for predicting patient survival, both in treated and treatment-naive patients [12]–[18].

Traditionally, detection of somatic point mutations is done using whole exome or genome sequencing of tumor and matched-normal DNA [19]–[23]. The latter is required for distinguishing between somatic mutations found exclusively in the tumor sample, and germline variants shared by all cells of an individual. Recently, several studies have developed a ’tumor-only’ pipeline that uses DNA sequencing to detect somatic mutations without the matched-normal sample, at the cost of lower precision and recall [24]–[26]. An additional extension to these pipelines includes the detection of somatic mutations from RNA sequencing and a matched-normal DNA sample. In a recent publication we have introduced such a pipeline termed RNA-MuTect, and showed that most of the mutations detected only in the RNA are filtered out by our pipeline, achieving an overall high precision. In addition, high sensitivity for mutations with sufficient detection power was observed, enabling the detection of most driver genes and mutational signatures [27].

In this study we took our RNA-based approach one step further and developed a pipeline for detecting somatic point mutations from RNA sequencing without a matched-normal sample, named RNA-MuTect-WMN (WMN; without-matched-normal). This is accomplished via a machine learning framework which utilizes a few dozens of features to classify single nucleotide variants as either somatic or germline. Our pipeline is trained and tested on the TCGA melanoma dataset where it achieves high precision and recall. In addition, it enables a reliable identification of both driver genes and mutational signatures. When applied to estimate the TMB from RNA samples alone, we find that its performance is either equivalent or superior to TMB estimated based on DNA with a matched-normal sample. The ability to estimate the TMB using a single RNA sample further emphasizes the potential clinical utility of our pipeline.

## Results

### Identifying somatic mutations from RNA-seq data without a matched normal sample

To develop a pipeline for detection of somatic point mutations from RNA-seq without a matched-normal sample, we leveraged RNA-seq and matched-normal DNA of 462 melanoma samples from The Cancer Genome Atlas (TCGA) [28]. To obtain the ground truth of somatic and germline variants in these samples, we ran RNA-MuTect [27]; in short, RNA-MuTect works by first running MuTect [29] on tumor RNA and matched-normal DNA, to identify the set of tumor somatic mutations and the set of potential germline variants (Methods). Since this set includes multiple noisy sites unique to RNA, a series of filtering steps is then applied to yield the final set of true somatic mutations (Figure 1A). To examine the accuracy of RNA-MuTect on this set of samples, we compared the list of somatic mutations to that obtained using tumor and matched-normal DNA. As originally reported [27], focusing on the RNA mutations with sufficient detection power in the DNA, 90% were indeed found in the DNA, with a median of only 3 detected mutations per sample found in the RNA alone (Methods).

**Figure 1:**
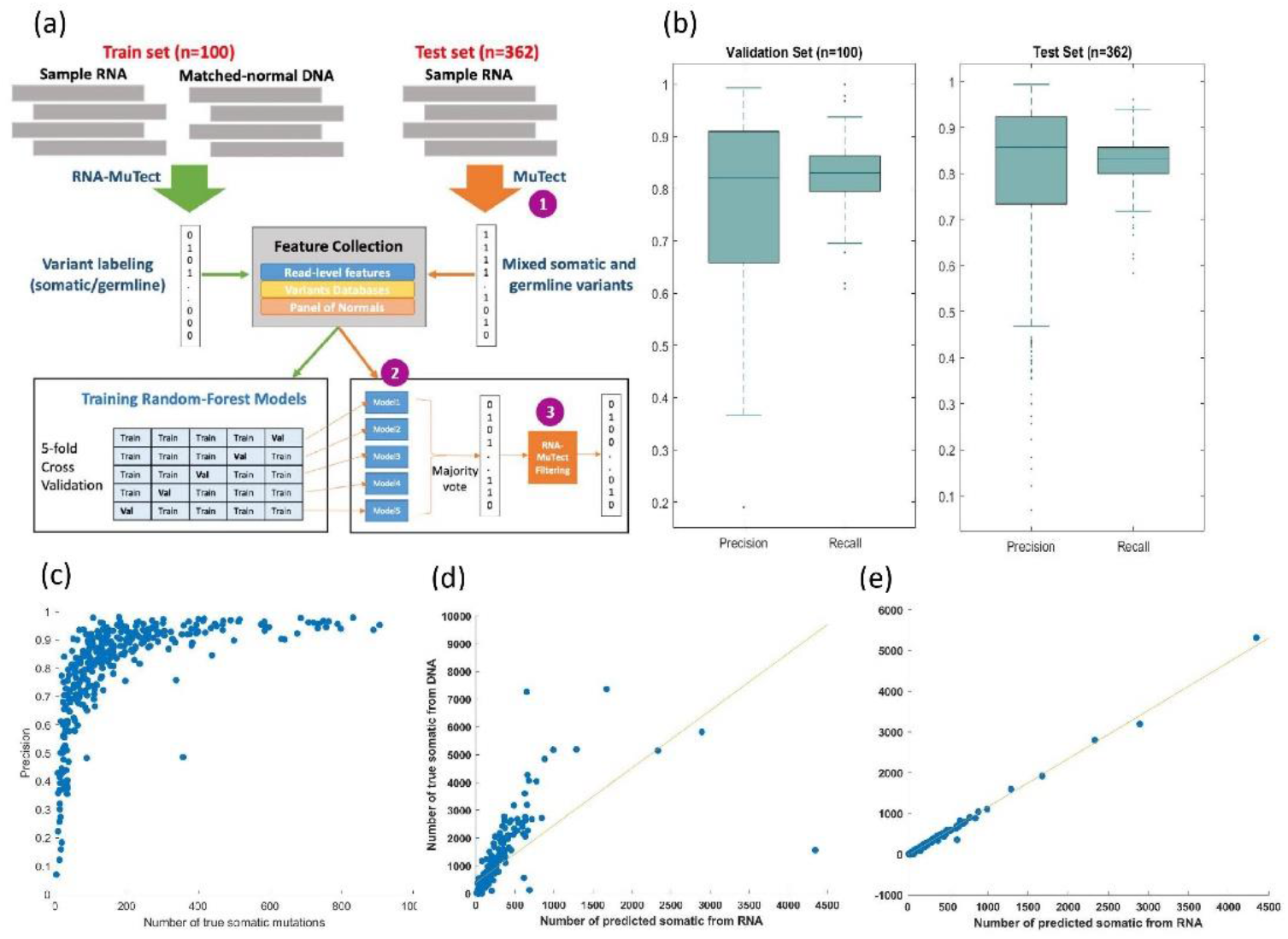
(a) An overview of the RNA-MuTect-WMN pipeline: In the training set (n=100, green arrows), RNA-MuTect is applied on tumor RNA and matched-normal DNA to obtain a list of variants labeled as somatic or germline. A random forest classifier is then trained with the collected set of features for each variant in a 5-fold cross validation manner. In the test set (n=362, orange arrows), (1) MuTect is applied with tumor RNA and without a matched-normal sample, to yield a list of mixed somatic and germline variants. (2) The five trained models are then applied to this set of variants and classifies them as either somatic or germline in a majority vote manner. (3) Finally, the predicted set of variants is further filtered by the RNA-MuTect filtering steps. (b) Precision and recall on validation (left) and test (right) sets computed for each sample. Box plots show median, 25th, and 75th percentiles. The whiskers extend to the most extreme data points not considered outliers, and the outliers are represented as dots. (c) Precision as the function of the number of true somatic mutations per sample. (d) Correlation between the number of predicted somatic mutations and the number of somatic mutations as determined by DNA with matched-normal DNA sample. (e) Correlation between the number of predicted somatic mutations and the number of somatic mutations as determined by RNA with a matched-normal DNA sample.

To classify point mutations as either somatic or germline, we collected for each variant a set of genomic features (Methods). This list includes the number of reference and alternate reads, variant classification type and MuTect likelihood score. In addition, we collected data on germline variants from dbSNP [30], gnomAD [31], 1000 genomes [32] and the Exome Sequencing project [33]. Finally, we utilized both DNA and RNA panel of normal (PoN) which are based on ∼8000 TCGA and ∼6500 Genotype-Tissue Expression (GTEx) normal samples (Methods) [34]. These PoNs encode the distribution of alternate read counts across the entire sets of normal samples [35].

To test how well our features separate between somatic and germline variants, we performed a Wilcoxon rank sum test for each feature, and found that all features show a significant difference between these two types of variants (FDR corrected p-values <= 0.0111, Supplementary Figure 1). However, when searching across a range of thresholds in each feature, we found that the Precision-Recall Area Under the Curve (PR-AUC) is very low (<0.08, Supplementary Figure 2), as well as the F1 score (<0.16, Supplementary Figure 3). This finding is a result of the substantial overlap between features’ values in these two variant types, demonstrating the need for a more complex model.

To this end we developed a machine learning framework named RNA-MuTect-WMN that gets as input a list of variants with their associated features, and classifies them as either somatic or germline variants as follows: first, our data is randomly split into training (n=100) and test sets (n=362). In the training set, each sample contains a list of single nucleotide variants with their genomic features (Methods), and a somatic\germline label based on the RNA-MuTect pipeline, as described above (Figure 1a). Next, a set of random forest classifiers is trained [36] in a 5-fold cross validation manner, such that in each iteration 80 samples are used for training and 20 samples are used for validation. The median precision and recall achieved by our model when computed on each sample in the validation set are 0.82 and 0.83, respectively (Figure 1b). To test our model performance we used our test set of 362 samples and applied the following three steps: (1) we ran MuTect with tumor RNA-seq and *without* a matched-normal sample. In this step both somatic and germline variants are marked as true somatic mutations, and a subset of sites are removed based on MuTect filtering scheme; (2) we then applied the 5 models built in the training step, and classified each variant as either somatic or germline based on the majority vote; (3) finally, to remove any remaining RNA-specific noise, we applied the RNA-MuTect filtering steps and achieved the final set of predicted somatic mutations. We have decided to run RNA-MuTect filtering steps on the narrowed list achieved after step 2 instead of upfront at step 1, due to a couple of time consuming steps implemented in the pipeline that could have significantly slow down the process. The final set of somatic and germline variants was then used to estimate the pipeline’s performance, showing a sample-level median precision and recall of 0.85 and 0.83, respectively (Figure 1b).

Further investigating our results, we observed that a few samples achieved a relatively low precision (41 out of 46 samples with precision < 0.6 had mutation count < 50, Figure 1c). We found that this performance is due to the overall low number of somatic mutations in these samples, and that the median precision on the remaining samples is 0.89. In addition, to circumvent the possibility that the high performance obtained by our model is a result of low purity levels which will in turn result with substantially different allele fractions for somatic and germline variants, we examined the correlation between tumor purity and the obtained precision and recall levels. Encouragingly, we found this correlation to be insignificant (Spearman R = -0.0040, -0.0874, P-value = 0.93, 0.09, for precision and recall, respectively).

To better characterize our model we next examined which features are the most important in distinguishing between somatic and germline variants, using the feature importance score (Methods). We found that a few of the PoN features as well as the gnomAD feature are the most influential in our model (Supplementary Table 1). Finally, we computed the Spearman correlation between the number of predicted somatic mutations and the number achieved by the DNA or RNA with a matched-normal DNA sample. In both cases, we found it to be highly significant (R=0.92, P-value = 4.15^-151^ for DNA and R = 0.98, P-value < 8.7*10^−286^ for RNA, Figure 1d-e, respectively).

### Detecting mutational signatures and significantly mutated genes without a matched-normal sample

The overall high performance of RNA-MuTect-WMN enabled us to apply our standard analysis pipelines for identifying mutational signatures and significantly mutated genes. To this end we applied SignatureAnalyzer [37], [38] using the set of predicted somatic mutations, and identified 4 signatures (Figure 2a): UV signature (COSMIC SBS7b, cosine similarity = 0.95) which is common in melanoma [39], [40], signature 5 (COSMIC SBS5, cosine similarity = 0.87) which is common in various cancer types, including melanoma, and a signature enriched with C>A mutations that was previously found in ultraviolet light associated melanomas (SBS38, cosine similarity = 0.78). Importantly, the same three signatures were identified in the DNA (Supplementary Table 2 and Supplementary Figure 4). In addition, a signature enriched with T>G mutations was detected. This signature was not detected in the DNA but was detected in the RNA when somatic mutations were identified with a matched-normal DNA sample (Supplementary Figure 4). Indeed, we found that out of 552 mutations that are associated with this signature, 489 were detected only in the RNA. While it is hard to conclude whether this signature is a true RNA signature or a result of RNA-specific noise, it is important to note that its detection is not specific to our pipeline in which a matched-normal sample is not used.

**Figure 2:**
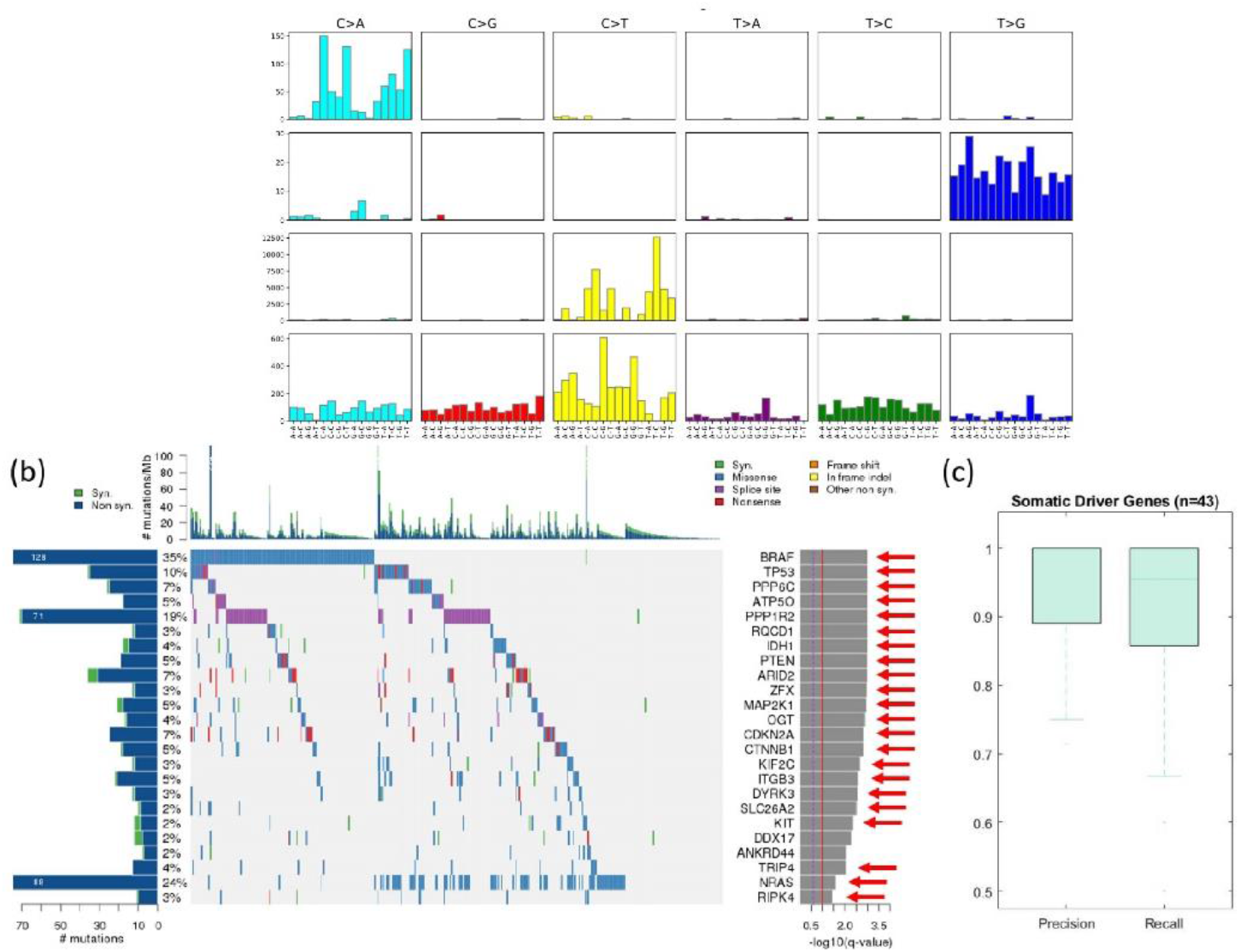
(a) Mutational signatures identified by SignatureAnalyzer [37], [38] on the basis of predicted somatic mutations; (b) Co-mutation plot based on predicted somatic mutations in our test set. Overall frequencies, allele fractions, and significance levels of candidate cancer genes (Q < 0.05) identified by MutSig2C [41] are shown. Genes marked with a red arrow were also identified as significantly mutated based on the set of somatic mutations detected using RNA and a matched-normal DNA. (c) Precision and recall on the set of know melanoma drivers. Box plots show median, 25th, and 75th percentiles. The whiskers extend to the most extreme data points not considered outliers, and the outliers are represented as dots.

Next, we identified significantly mutated genes by applying MutSig2CV [41] on the set of predicted somatic mutations. Out of 24 identified genes, 22 were found to be significantly mutated also when the matched-normal sample is taken into account (Figure 2b), and only 2 were missed by our pipeline (Supplementary Table 3). Importantly, 15 out of the 24 genes were also identified as significantly mutated based on a DNA analysis, a rate that is similar to our previous report [27]. Finally, we examined our pipeline’s performance in identifying a set of 55 known melanoma somatic driver genes found in the COSMIC database [42] (Supplementary Table 4). We found that on the set of 43 genes that carried at least one true somatic mutation in our dataset, our pipeline achieves an even higher precision and recall, with median values of 1 and 0.95, respectively, further demonstrating its high value.

### TMB predicted by RNA-MuTect-WMN is associated with patient survival

The development of checkpoint blockade (CPB) therapy such as anti-PD1 and anti-CTLA4 has revolutionized cancer therapy and resulted in long-lasting tumor responses in patients with a variety of cancers [43]. As a result, these drugs have been FDA-approved for many cancer types, including melanoma, non-small cell lung cancer, Urothelial carcinoma, Head and Neck squamous cell carcinoma and more [44]. Recently, an accelerated approval for anti-PD1 for the treatment of adult and pediatric with tumor mutational burden-high (TMB-H, ≥ 10 mut/Mb) has been granted, making it a critical metric in the clinical decision process. Indeed, the TMB which is traditionally estimated via DNA sequencing, has been found to be associated with patient survival to different extents, depending on cancer type [45], prior and current treatment [46]–[48].

Here, based on the set of predicted somatic mutations from RNA sequencing alone, we estimated the TMB as the number of non-silent somatic mutation in each sample. We then divided the patients into two groups with high- and low-TMB levels, using the median TMB as the cutoff value. We found that patients with high-TMB had a mild but significant increase in survival time as compared to those with low-TMB (log-rank P-value = 0.02, Figure 3a). Of note, performing the same analysis using the set of somatic mutations detected based on tumor and matched-normal DNA, similar results are obtained (logrank P-value = 0.01, Figure 3b), further demonstrating the utility of our pipeline.

**Figure 3:**
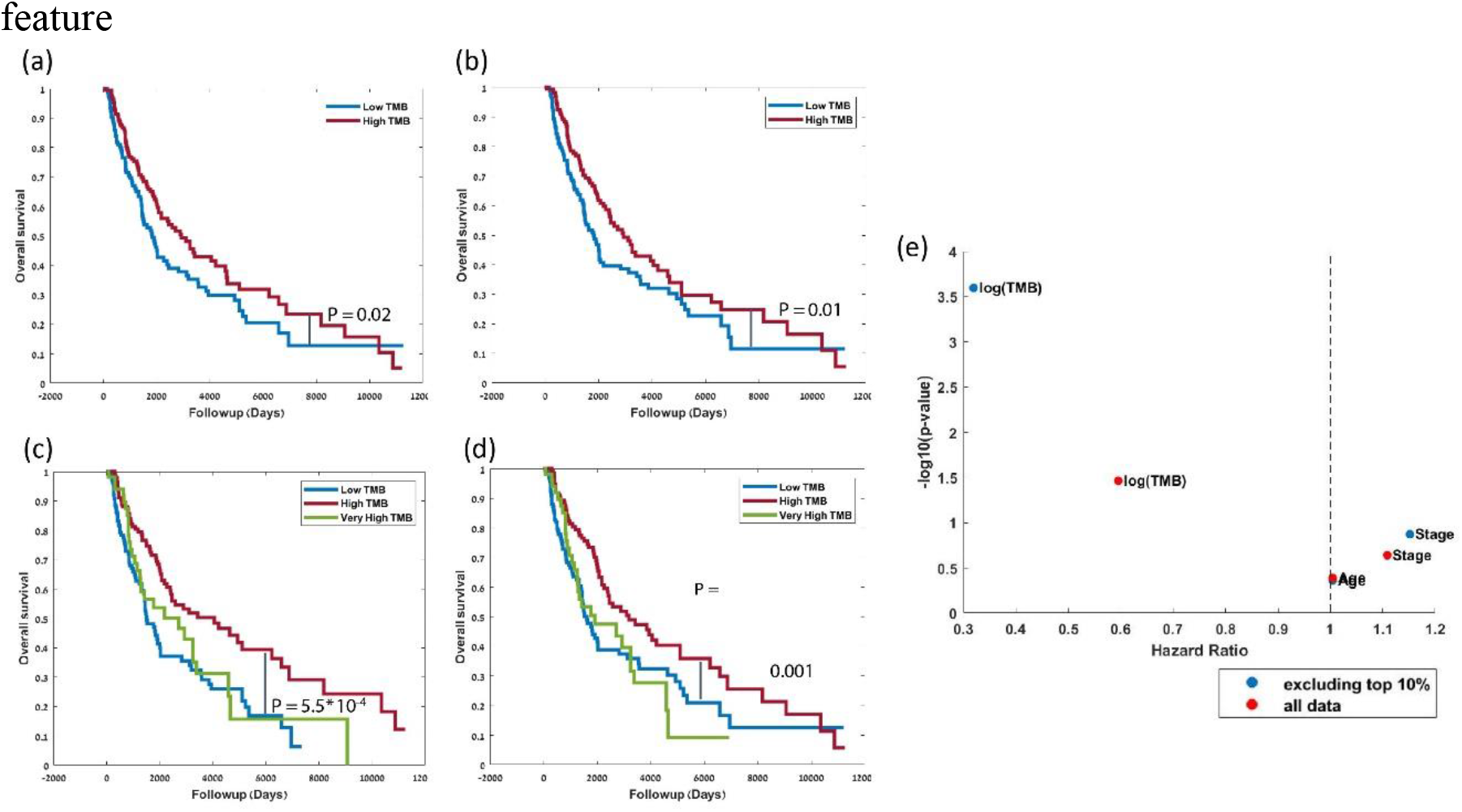
(a-b) Kaplan–Meier survival curves for patients with high vs. low TMB estimated on the basis of predicted somatic mutations from RNA alone (a); or on the basis of DNA with a matched-normal sample (b). The median TMB value is used to define the ‘low TMB’ and ‘high TMB’ subgroups. The P-value is computed using a log-rank test. (c-d) Kaplan–Meier survival curves as in (a) and (b), respectively, with patients divided into three groups with very-high vs. high vs. low TMB. Subgroups were split by using the top 10^th^ percentile for the very high group, and median for the remaining samples. (e) Hazard Ratio vs. –log_10_(p-value), obtained by a multivariate Cox proportional hazards regression analysis. Red dots represent the values obtained when all samples are used and blue dots represent the values obtained after excluding the top 10% of samples (very high TMB).

Performing a multivariate Cox proportional hazards regression analysis with patient age, tumor stage and our RNA-based TMB estimates as the covariates, we found that TMB is the prognostic factor most associated with increased survival (HR = 0.59, 95% CI=0.36-0.96, P-value < 0.03, Figure 3e).

The extent of association between TMB and patient survival vary widely between the different datasets according to cancer type and prior therapy. A recent publication by Valero et al. showed that among patients that were not treated with CPB, a very high TMB at the top percentiles is associated with poor survival [49]. Given that most of the patients in the TCGA cohort were not treated with CPB, we set to examine this observation in our data as well. Indeed, when we divide the patients into three groups with very high-, high- and low-TMB levels, using the top 10^th^ percentile for the very high group, and median for the remaining samples, we find that those with the highest TMB values have a poor survival (logrank P-value = 0.04 between high- and very high-TMB), and those with median high TMB have an improved survival as compared to those with low TMB (logrank p-value = 5.8*10^−4^, Figure 3c). This result is robust to the selection of threshold for the very-high TMB group (top percentile between 10^th^ -15^th^). Importantly, performing the same analysis based on DNA revealed the same trends, though with an inferior significance level (logrank P-value = 0.01, 0.04, respectively, Figure 3d). Repeating the Cox regression analysis while removing the top 10^th^ percentile, the association of TMB with survival became more significant (HR = 0.31, 95% CI=0.17-0.58, P-value < 2*10^−4^, Figure 3e).

Overall, these results demonstrate that estimating TMB based on RNA alone is feasible and of a high predictive power.

### An improved RNA-based TMB estimation in patients treated with CPB

We next examined the prediction power of our model on an additional set of patients that were treated with nivolumab (anti-PD1), some were treatment naive and some had previously progressed on ipilimumab (anti-CTLA4) [50]. Raw RNA-sequencing data from 50 pre-therapy biopsies were obtained and aligned to the reference genome (Methods). Then, RNA-MuTect-WMN was applied to identify and classify the set of somatic mutations in each sample. To validate our calls, we first applied SignatureAnalyzer and identified the set of mutational signatures that are active in these samples. Encouragingly, we found the UV signature (SBS7b), along with the TMZ signature (SBS11) and SBS5 that were also found by the authors based on DNA (cosine similarity = 0.86, 0.95 and 0.78, respectively, Figure 4a). In addition, when applying MutSig2CV to identify significantly mutated genes, both NRAS and BRAF, known melanoma drivers, were found to be significantly mutated (Figure 4b).

**Figure 4:**
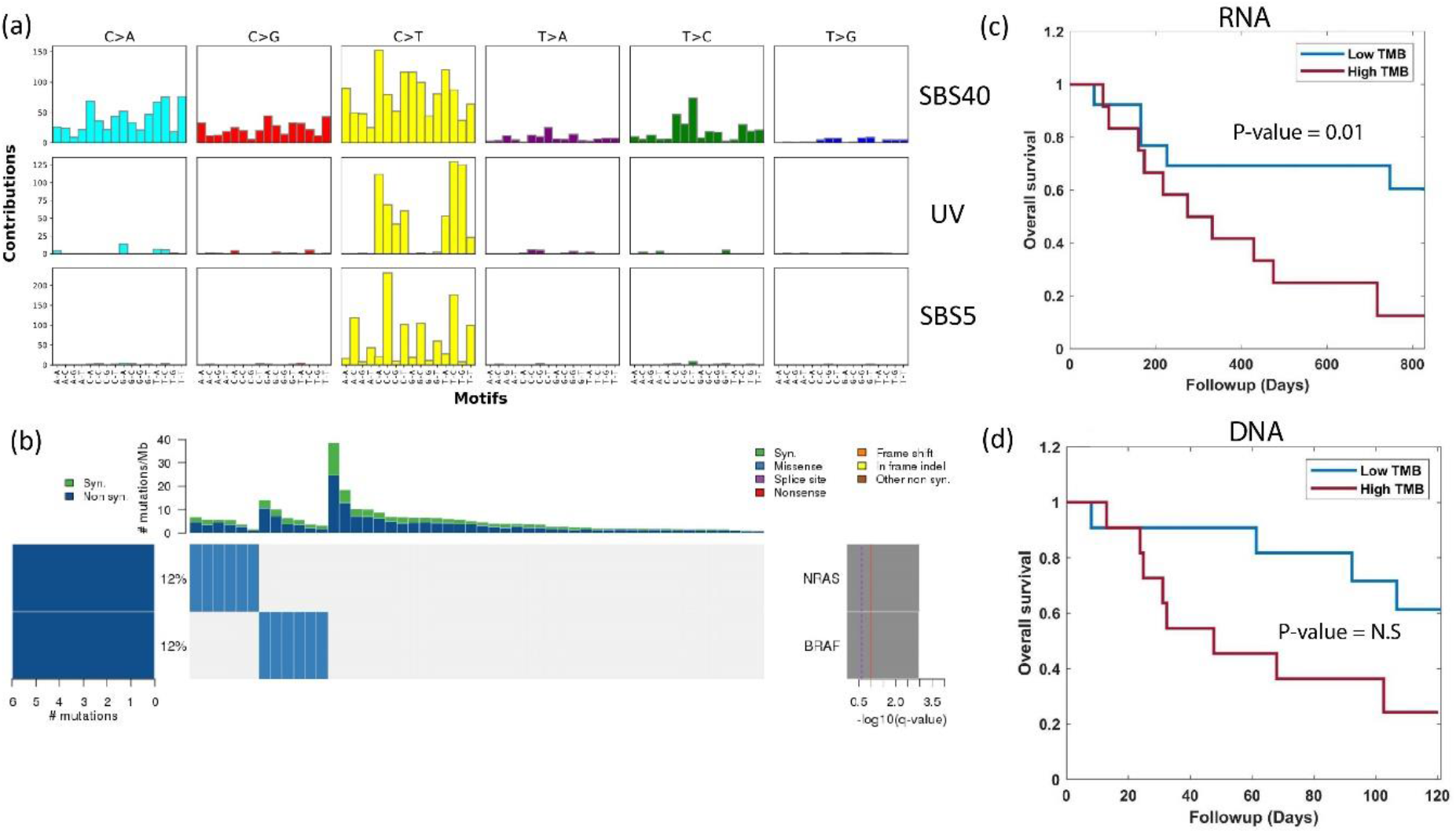
(a) Mutational signatures identified based on the set of predicted somatic mutations using the RNA-seq data of 50 pre-therapy biopsies from the Riaz et al. [47] dataset. (b) Co-mutation plot based on predicted set of somatic mutations. Overall frequencies, allele fractions, and significance levels of candidate cancer genes identified by MutSig2CV [41] are shown. (c-d) Kaplan–Meier survival curves for patients that have previously progressed on ipilimumab. TMB is estimated based on predicted somatic mutations from RNA and based on the list of somatic mutations detected by the authors using tumor and matched-normal DNA (d). The patients are split to two groups of ‘low TMB’ and ‘high TMB’ using the median TMB value as the cutoff.

Finally, we estimated the TMB based on the set of predicted somatic mutation. Interestingly, when considering the set of treatment naive patients for which both DNA and RNA are available, no significant association between TMB and patient survival is found, based on neither DNA nor RNA. However, when considering the set of patients that were previously progressed on ipilimumab, a significant association between high TMB and poor survival is found (logrank P-value = 0.01, Figure 4c). This is in similar to the trend reported by the authors using DNA (Figure 4d), which was insignificant. Overall, in this independent dataset as well we find that estimating the TMB from tumor RNA alone is feasible and results with similar trends to those obtained with tumor and matched-normal DNA.

## Discussion

In this study we introduce RNA-MuTect-WMN, the first computational method that identifies somatic mutations from RNA-seq data without a matched-normal sample. Our pipeline is based on the RNA-MuTect method [27] which is designed to detect somatic mutations from tumor RNA-seq and matched-normal DNA. To extend it to a ’tumor-only’ mode we built a random forest classification model that distinguishes between somatic and germline variants using various features, including both mutation specific ones and those derived from large panels and databases of normal individuals. Our model was trained on a subset of the TCGA melanoma dataset, and achieved high precision and recall (85% and 83%, respectively) when applied to an independent test set of additional >350 melanoma samples. Importantly, we find that when using the set of predicted somatic mutations which are derived from RNA-seq samples alone, all mutational signature and >90% of the driver genes are correctly identified, as compared to RNA-MuTect, where a matched-normal sample is used.

When calling somatic mutations directly from RNA-seq we are clearly limited to the set of mutations that are sufficiently expressed. While this reduces sensitivity for certain downstream analyses, it may increase it in others. Specifically, we hypothesized that estimating the tumor mutational burden from RNA rather than from DNA would improve prediction power, as only expressed mutations can become neoantigens and elicit an immune response. Indeed, we first show that estimating TMB from tumor RNA-seq without a matched-normal sample is feasible, and that the exact same trends as those estimated using tumor DNA with a matched-normal sample are observed. Moreover, the prediction power of RNA-based TMB is either equivalent or superior to that estimated by DNA. As previously shown, we find that in melanoma patients that were not treated with CPB, very high TMB is associated with poor survival [49], while median high is associated with improved survival as compared to patients with low TMB. In addition, in treated patients that were previously progressed on anti-CTLA4, we find that high TMB is significantly associated with poor survival compared to low TMB. These results are in concordance with the original findings [15].

Importantly, the model built in this study is based on melanoma samples, and preliminary results show that a different model is required in order to achieve a high performance in other cancer types (data not shown). However, the RNA-MuTect-WMN approach is generic and can be easily applied for any cancer type, given a sufficient number of samples with RNA-seq of the tumor, along with tumor and matched normal DNA for validation. Moreover, melanoma is a highly mutated cancer with a sufficient number of somatic mutations that can be used for model training, and where the fraction of germline contamination predicted by our model is negligible. Hence, the performance of our approach should be further tested on lowly mutated cancers where significantly less somatic mutations are available for training, and where the fraction of germline contamination can become substantial. These limitations can be potentially addressed by down-sampling of the germline group, or by combining multiple datasets together.

Overall, we believe that the motivation for using RNA-MuTect-WMN is three-fold: first, for future studies, it diminishes the need for collecting and sequencing matched-normal samples, thus significantly reducing sequencing cost, especially for large cohort analysis. Second, it enables the analysis of RNA-seq data in retrospective studies where RNA was originally sequenced for expression-based analyses. Third, it enables a combined analysis where both genetic and phenotypic data can be inferred from the exact same sample. This is especially crucial in cancer where different regions of a tumor from which DNA and RNA are extracted may be significantly different due to tumor heterogeneity. These applications can significantly increase the number of samples analyzed and thus aid biomarker and drug target discovery.

## Methods

### DNA Mutation calling pipeline

TCGA DNA BAM files aligned to the NCBI Human Reference Genome Build GRCh37 (hg19). Sample contamination by DNA originating from a different individual was assessed using ContEst [51]. Somatic single nucleotide variations (sSNVs) were then detected using MuTect [52]. Following this standard procedure, we filtered sSNVs by: (1) removing potential DNA oxidation artifacts [53]; (2) realigning identified sSNVs with NovoAlign (www.novocraft.com) and performing an additional iteration of MuTect with the newly aligned BAM files; and (3) removing technology- and site-specific artifacts using a panel of ∼8000 TCGA normal samples (PoN filtering, see [54]). Finally, sSNVs were annotated using Oncotator [55].

### RNA mutation calling pipeline (RNA-MuTect)

RNA FASTQ files were downloaded from the Genomic Data Commons database aligned to the NCBI Human Reference Genome Build GRCh37 (hg19) using STAR [56]. The RNA-MuTect pipeline was then applied as previously described [27]. In short, MuTect is first applied using the ALLOW_N_CIGAR_READS flag, with a STAR-aligned RNA-seq BAM and a matched-normal DNA-seq BAM. The first set of somatic mutations is then filtered with a DNA PoN as described in the ’DNA Mutation calling pipeline’ section. A series of filtering steps is then applied, including a realignment step with HiSat2, a RNA PoN based on GTEx samples, removal of RNA editing sites and more, as previously described in detail [27].

### Power analysis

Given a mutation with an alternate allele count of *x* and a reference allele count of *y* in the RNA, we computed the power to detect it given a coverage *N* in the DNA. This was done by applying a beta-binomial model for observing at least *k* reads: 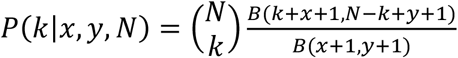 where *B* is the Beta function. To determine the minimal number of reads *k*, we first computed the error rate at the variant site, *r*, using the matched normal sample by taking the maximal allele fraction of the three possible alternate alleles and applying the Laplace correction with α=1. We then identified *k* as the number of alternate reads that have a probability <1% to be generated by the noise. Eventually, powered mutations were considered as those with power > 0.95 and alternate read count >=4.

### The RNA-MuTect-WMN pipeline

#### Data preprocessing

For the training set where a matched-normal sample is used, we labeled the set of variants that passed our entire calling pipeline as described in the ’RNA Mutation calling pipeline’ section as somatic. Germline variants were determined based on MuTect annotation as ‘normal_lod’, ‘germline_risk’ or ‘alt_allele_in_normal’. The analysis is focused only on chromosome 1-22, X and Y.

#### Feature collection

the following features were used in our pipeline:

1. T_ref_count - number of tumor reads supporting the reference allele
2. T_alt_count - number of tumor reads supporting the alternate allele
3. T_lod_fstar – The LOD score computed by MuTect
4. Tumor_f – tumor allele fraction

5-12. For each of the germline variants database (dbSNP, gnomAD, 1000Genome, ESP) two vectors were created: (a) A Boolean indicating whether the variant is present (1) or not (0) in each database; (b) variant allele fraction (AF), when available, and a mean AF value over all variants in the database when this data is missing.
13. Variant_classification - if the variant classification as defined by Oncotator [57] was either IGR, Intron, RNA, lincRNA this feature was set to be (1) and (0) otherwise. We included this feature following an analysis showing that almost no somatic mutations were annotated with this variant classification.
14-32. DNA and RNA Panel of Normals –each genomic position in each PoN is binned into one of eight bins using its allele fraction as previously described [35]:

1. total counts < 8 (insufficient coverage)
2. total counts >= 8 (and no alternate reads above subsequent thresholds)
3. alt count >= 1 and alt fraction >= 0.1%
4. alt count >= 2 and alt fraction >= 0.3%
5. alt count >= 3 and alt fraction >= 1%
6. alt count >= 3 and alt fraction >= 3%
7. alt count >= 3 and alt fraction > 20%
8. alt count >= 10 and alt fraction >= 20%

The 9^th^ feature for each PoN is then the log likelihood score computed as in [35].

#### Model training

100 samples were randomly selected and defined as the training set. These samples were then divided to 5 pairs of training and validation sets with 80 and 20 samples in each group, respectively. A random forest classifier was applied on each of the training sets, using the somatic and germline labels. Each resulting model was then tested on the corresponding validation set, and the precision and recall were calculated to evaluate the performance of all models.

#### Model testing

To test the models generated in the previous step we first applied MuTect on our test set composed of the remaining 362 samples, using tumor RNA-seq and without the matched normal sample. As a result, we obtained a list of variants containing both somatic, germline and RNA-specific noise. For each of these variants we collected the set of features described above and applied the 5 trained models. Each variant was then assigned a somatic or germline label based on a majority vote of the 5 models. Finally, the predicted group of somatic mutations was further filtered using RNA-MuTect filtering steps, as described in [27].

### Signature analysis

To identify mutational signatures, we used the SignatureAnalyzer tool: https://github.com/broadinstitute/getzlab-SignatureAnalyzer [38]. A cosine similarity score was used as a measure of closeness to known signatures. This score ranges between zero and one, where a similarity of one represents identical signatures and a similarity of zero completely different mutational signatures. The similarity was measured against the latest version (V3.2) of SBS signatures in COSMIC.

### MutSig2CV for RNA-seq data

To apply MutSig2CV [41] for RNA-seq data, we utilized gene-level coverage model that reflected which bases were typically sufficiently covered in each gene using GTEx RNA-seq data, as previously done [27]. We considered genes as significantly mutated if they had an FDR-corrected *Q* value < 0.05.

### Riaz data analysis

Raw RNA sequencing data for 50 available pre-treatment samples was aligned to the NCBI Human Reference Genome Build GRCh37 (hg19) using STAR [58]. As described in the main text (Figure 1a), we then ran the three steps of our pipeline: (1) running MuTect with tumor RNA alone; (2) applying the 5 models trained on TCGA data to get an initial list of predicted somatic mutations; (3) applying RNA-MuTect filtering steps. The final list of somatic variants was used for identifying significantly mutated genes, mutational signatures and for estimating the TMB.

### Tumor Mutational Burden Analysis

To compute tumor mutation burden (TMB), we counted the number of non-silent somatic SNVs per sample. For the survival analysis, the absolute TMB value was used for determining the median TMB and the top 10-15 percentile values. For the continuous Cox regression models we used log_10_(TMB) together with patient age and tumor stage. A logrank test was used to estimate the significance level of the survival analysis.

### Feature Importance

To calculate the feature importance we used the built-in feature importance scores of scikit-learn, also known as GINI importance (or-mean decreased impurity). We obtained the feature importance scores for each of the 5 trained models, and computed the final importance score for each feature based on the average score across all 5 models.

### Estimating precision/recall level for single features

For each feature, we computed the difference between germline and somatic variants using the Wilcoxon rank-sum test.

To calculate the best precision and recall that can be achieved, we computed the F-score across a range of thresholds and report the maximal one. The precision-recall AUC was computed in a standard way across a range of thresholds.

## Data availability

Access to TCGA raw DNA and RNA sequencing data (phs000178) was obtained via dbGap authorization. The Riaz [47] bulk RNA dataset used in this study is available under BioProject accession number PRJNA356761 or SRA SRP094781.

