## Supplementary material for "Estimating tumor mutational burden from RNA-sequencing without a matched-normal sample"

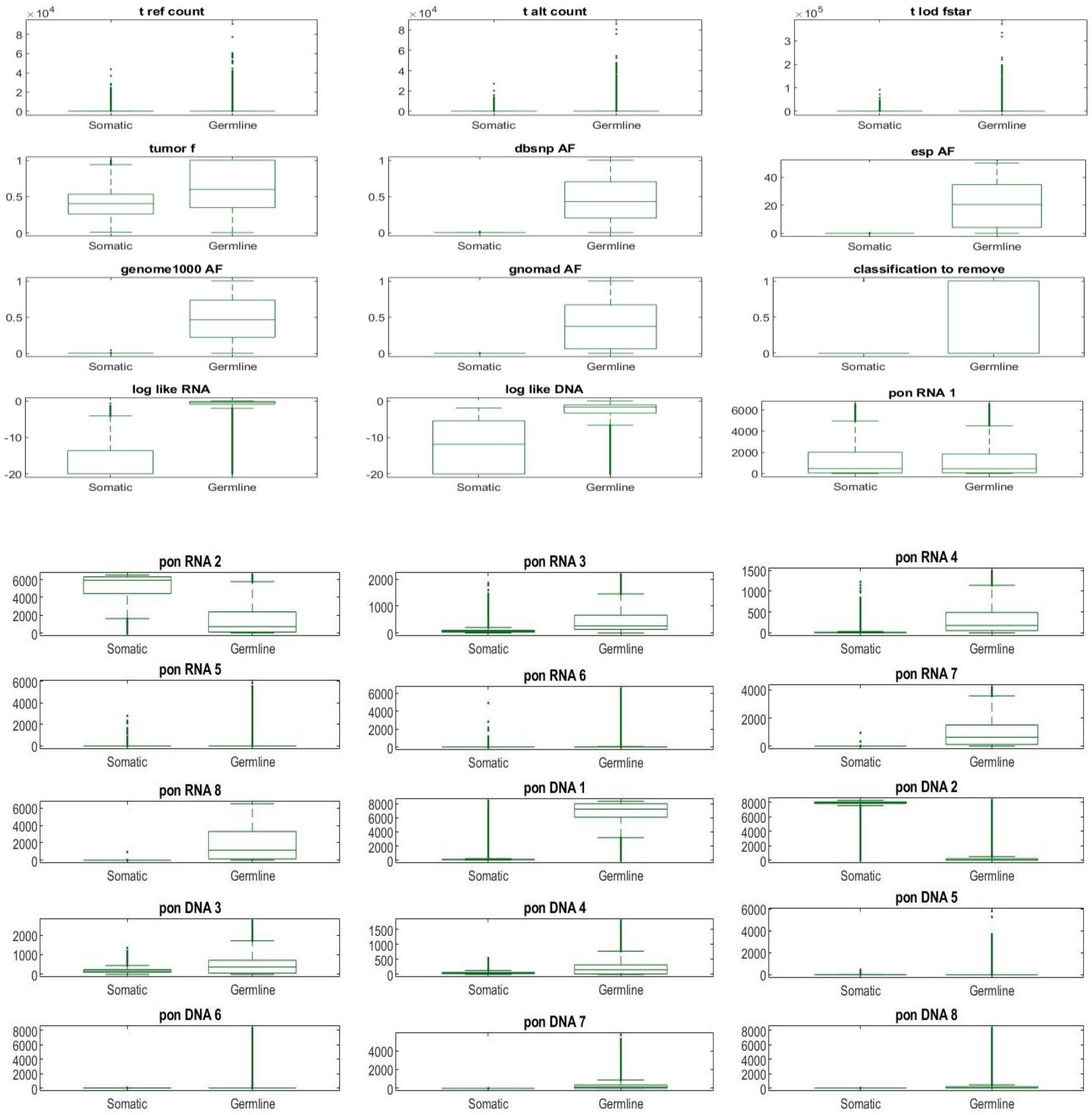


Supplementary Figure 1: Boxplots describing the different values between somatic and germline variants, in each feature separately.

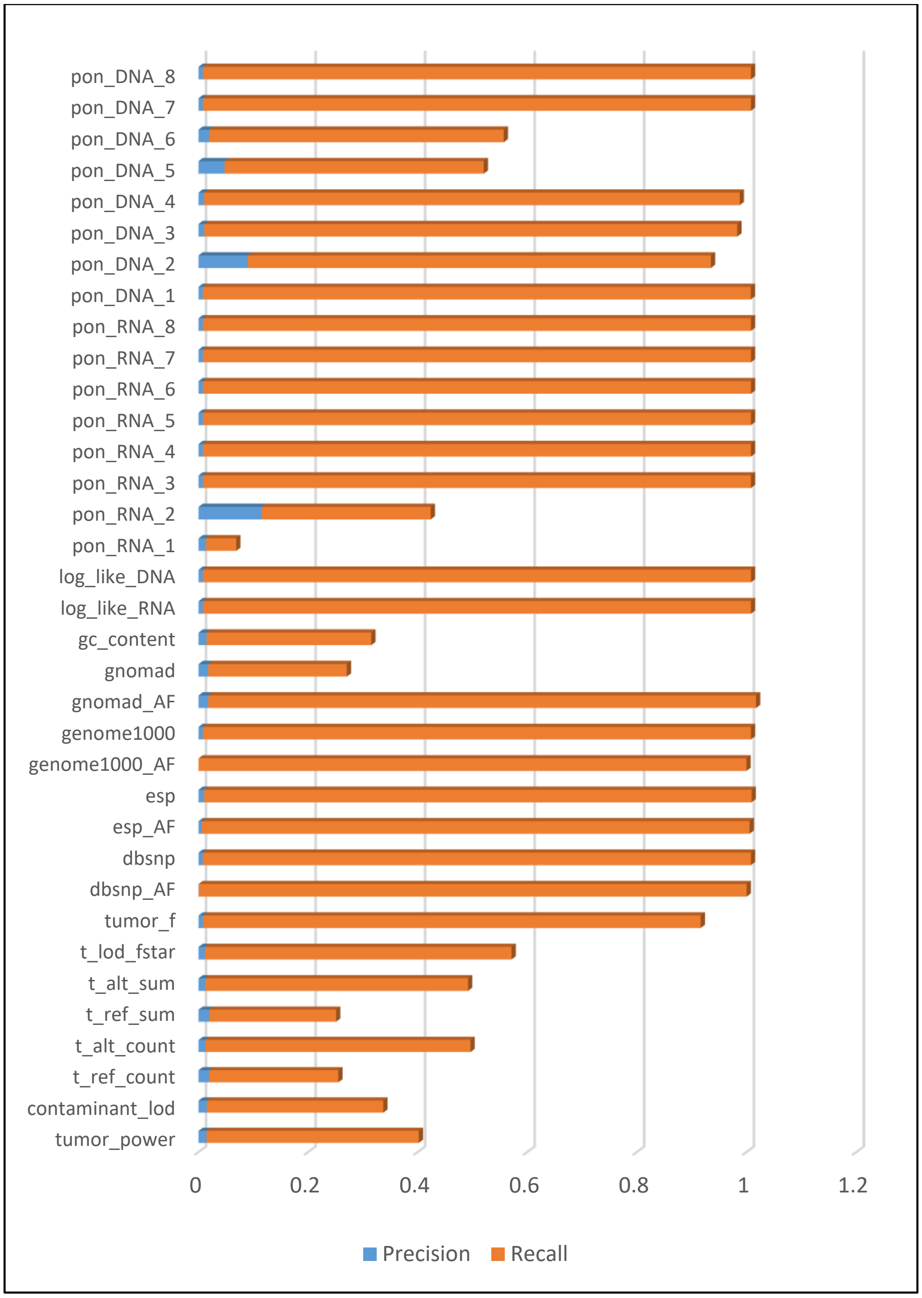


Supplementary Figure 2: Best precision and recall achieved for each feature, using the threshold providing the optimal F-score when calculated across a range of thresholds.

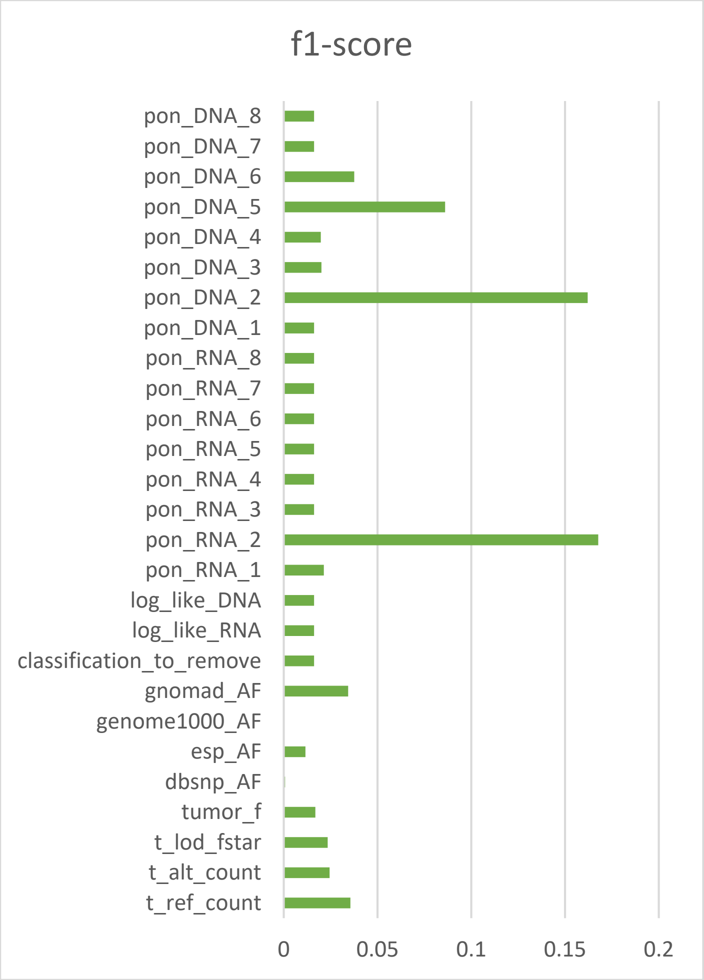


Supplementary Figure 3: Best F-score for each feature obtained by calculating the precision/recall across a range of thresholds.

(a)

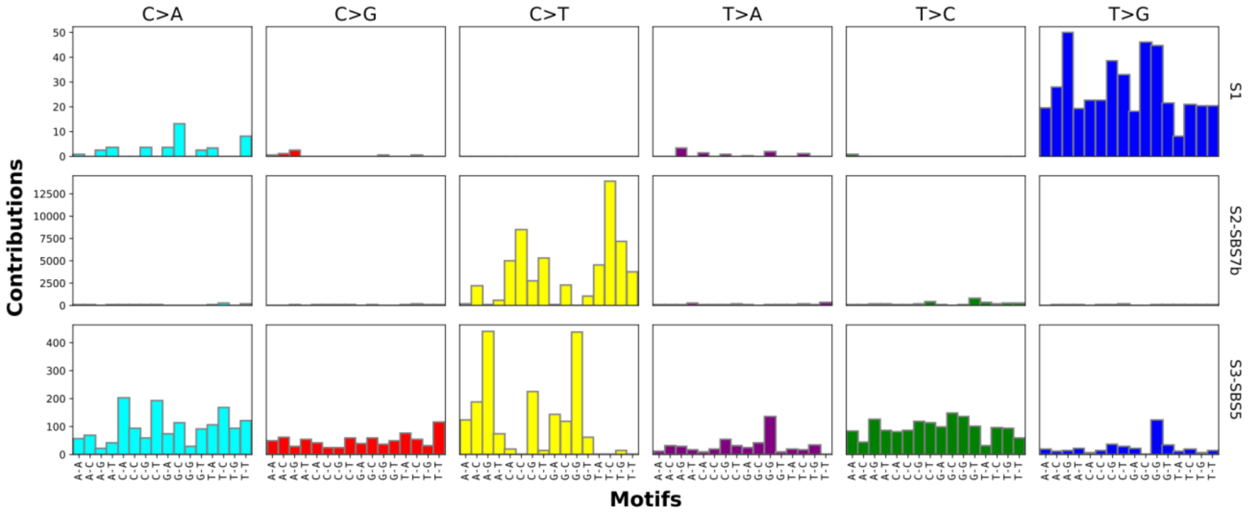


(b)

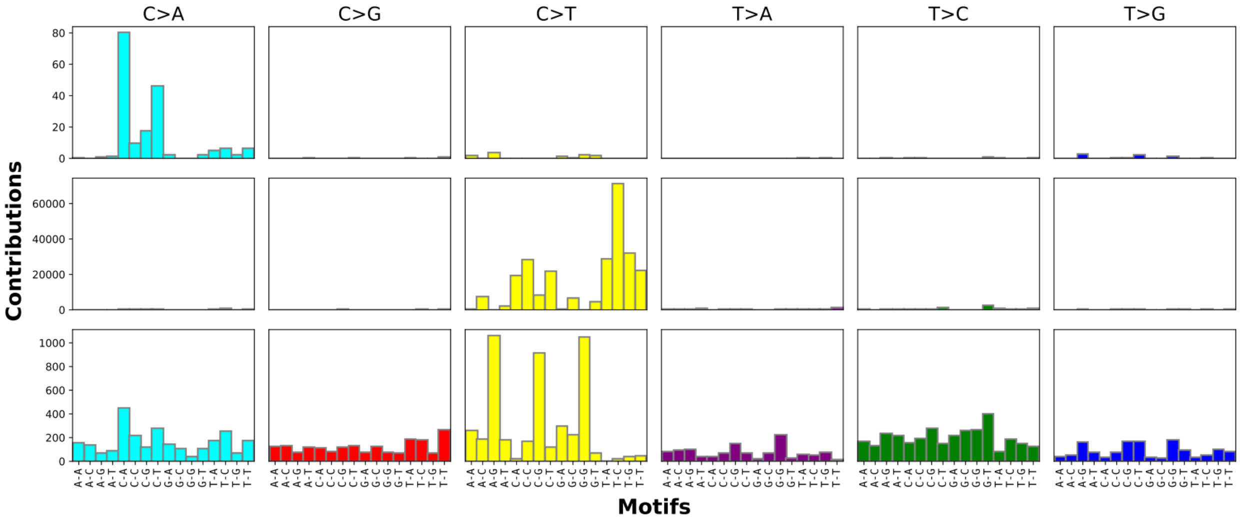


<u>Supplementary Figure 4:</u> Mutational signatures identified by SignatureAnalyzer [34], [35] on the basis of tumor RNA and matched-normal DNA (a) or on the basis of tumor and matched-normal DNA (b).
